# Topological Analysis of Vascular Networks: A Proof-of-Concept Study in Cerebral Angiography

**DOI:** 10.1101/2025.02.10.637397

**Authors:** Arturo Tozzi

## Abstract

The application of topological methods to cerebral angiography may provide a robust mathematical framework for analyzing cerebrovascular structures at multiple scales. In this proof-of concept study, we explored the use of algebraic and differential topology to characterize structural integrity, connectivity, flow dynamics and hierarchical organization of cerebral vascular networks. Through a hierarchical approach, we examined the topology from general to local, capturing macroscopic vascular organization down to individual vessel bifurcations. By leveraging key theorems, we assessed various aspects of topological analysis, including evaluation of total features, transition from total to local features, evaluation of local features, transition from local to total features, interaction between total and local features. These steps enable the analysis of the global connectivity of the vascular network, the detection of regional clusters and the identification of critical junctions at a local scale. A computational approach was developed to extract mathematical skeletons from angiographic images, constructing graph-based representations to study connectivity and homotopy equivalence. The Fourier decomposition of the vascular structures revealed dominant periodic patterns, indicative of structural stability and redundancy in the blood supply. Moreover, Betti number computations quantified vascular loops and branches, offering insights into collateral circulation potential. Our findings demonstrate that topological invariants can serve as diagnostic biomarkers for cerebrovascular diseases, including aneurysm susceptibility and ischemic risk assessment. This interdisciplinary methodology bridges mathematical topology with medical imaging, offering a novel lens for cerebrovascular analysis. Future work will integrate persistent homology and machine learning techniques for automated vascular topology classification.

## INTRODUCTION

Cerebral angiography plays a crucial role in diagnosing vascular disorders of the brain. Traditional methods rely on image-based assessments of vessel structure, flow dynamics, and morphological anomalies. However, these approaches often lack a formal mathematical framework to quantify connectivity and structural organization (Baharoglu et al., 2012; Kisler et al., 2017; Damseh et al., 2019; Ross et al., 2002; Li et al., 2021). Topological methods offer a powerful framework for analyzing vascular networks. Extending beyond conventional imaging techniques, they capture the intrinsic geometric and algebraic properties of the vasculature. Topology provides a means to analyze spaces independently of continuous deformations, making it particularly suitable for studying complex branching structures such as blood vessels (Bertolero and Bassett, 2010; De Domenico et al., 2015). A major challenge in angiographic imaging is the variability in vascular configurations among individuals, which can make traditional pattern recognition approaches difficult. Topological methods provide a framework for reducing this complexity by identifying fundamental properties of the vascular network that remain invariant under deformations. This proof-of-concept study applies a variety of topological theorems and computational methods to cerebral angiographic images to extract meaningful features, classify connectivity and detect possible vascular anomalies (Lauric et al., 2023; Gosh et al., 2024). By leveraging algebraic, differential, and computational topology, essential features of cerebral circulation can be captured, including connectivity, redundancy and flow optimization (He et al., 2008; Goirand et al., 2021).

Topological investigations of cerebral angiography may include global structural analysis, regional analysis, local vascular analysis, spectral methods. Global structural analysis of the vascular system mat be crucial for understanding blood supply distribution and redundancy. Using the Seifert-van Kampen theorem, we investigated how different connected components of the vascular network contribute to the fundamental group structure, providing insight into overall connectivity. Homology and Betti numbers allowed for the quantification of loops and independent vascular pathways, crucial in assessing the potential for collateral circulation in the event of arterial blockages. The interplay between arterial and venous structures was explored through the application of Poincaré duality, examining how these systems interact and preserve equilibrium in cerebral circulation.

Regional analysis focused on segmenting the vascular network into functionally relevant subregions, highlighting clusters that correspond to specific cerebral territories. The Künneth theorem facilitated this decomposition, allowing for homological computations across different vascular domains. Vascular segmentation may be particularly relevant in the study of ischemic stroke, where reduced perfusion in one territory can impact other regions. Graph-based methods further enhanced regional analysis by modeling the vasculature as a network of interconnected nodes and edges, where connectivity can be evaluated using group homomorphisms (Dummit and Foote, 2004). By leveraging the Whitehead theorem, we ensured that simplified graph representations preserve essential homotopy equivalence, retaining the critical structural features of the vascular system.

Local vascular analysis provided fine-grained insights into the branching patterns, bifurcations and junction points in the vascular system. Detecting bifurcation points and measuring their homological significance aided in identifying regions susceptible to stenosis, aneurysm formation or other pathological changes. The Hurewicz theorem helped in bridging homotopy and homology computations, facilitating the classification of vascular junctions. Knot theory, particularly Legendrian knot theory, was useful in detecting abnormal vascular loops or tangles that may indicate potential pathologies. The presence of specific knot structures in angiographic images may suggest hemodynamic stresses that may lead to vessel deformation or rupture.

Beyond static vascular analysis, spectral methods such as Fourier decomposition are employed to capture periodic and recurrent patterns in cerebral vasculature. By transforming angiographic images into frequency space, dominant structural patterns emerge, revealing how vascular pathways are organized at different scales. Inverse Fourier transforms allow for the reconstruction of dominant vascular features, filtering out noise and minor variations while retaining the essential geometry. This technique is particularly useful in differentiating normal vascular configurations from pathological formations, as deviations in frequency components can signal abnormal vessel growth or occlusion risks.

In sum, the integration of topological analysis with computational imaging may provide a robust framework for vascular assessment, moving beyond conventional image-based diagnostics. The combination of homotopy theory, homological computations and spectral analysis may enable a multi-scale understanding of cerebrovascular organization. By examining topology from a general perspective down to localized structural details, this proof-of-concept study aims to establish a methodology for quantifying vascular health and predicting potential disease risks.

## MATERIALS AND METHODS

We employed a series of computational techniques to analyze brain angiography images through topological and spectral methods, ensuring a comprehensive characterization of the vascular network. Our approach combined algebraic topology, graph theory, spectral analysis and image processing techniques to extract meaningful structural and functional information from the intricate vascular network. By leveraging these methods, researchers and clinicians can gain deeper insights into connectivity, continuity and spatial mapping of blood vessels. Our methodology was structured to progress from global analysis to regional segmentation and finally to local structural evaluation, allowing a hierarchical understanding of cerebrovascular connectivity. The following two paragraphs present: (a) a list of feasible theorems and concepts applicable to the topological analysis of brain angiography, and (b) a proof-of-concept experimental example demonstrating their application to real imaging data.

### Topological methods for analysing brain angiography

Various theorems evaluate different aspects of topological analysis, including the evaluation of total features, the transition from total to local features, the evaluation of local features, the transition from local to total features, the interaction between total and local features.

### Evaluation of total features

1. Borel’s theorem, which states that every sequence of independent random variables converges in probability (Borel 1953), may be used in probabilistic modeling of cerebral blood flow patterns. In brain angiography, understanding the distribution of contrast agents and their diffusion may be analyzed using this theorem, ensuring that variations in flow due to anatomical differences remain within predictable limits.
2. Coarse proximity theory may help in quantifying large-scale structures within the angiographic imaging data (Shi and Yao, 2024). By applying this concept, regions of vascularization can be compared across different subjects without being confounded by individual vessel structure variations. This may be particularly useful in stroke prediction and the study of large vessel occlusions, where macroscopic vascular topology plays a crucial role in determining collateral circulation.
3. Kolmogorov’s zero-one law, which deals with the behavior of tail events in probability spaces (Brzeźniak and Zastawniak, 2020), may predict whether vascular patterns lead to pathological conditions. If certain vessel formations or distributions occur with probability one, angiographic imaging data can be used to make deterministic predictions about disease development.

### Transition from total to local features

1. The Eilenberg-Zilber theorem, which describes the interaction between homology groups of spaces in product form (Golański and Lima Gonçalves, 1999), may be applied to the analysis of cerebral vessel connectivity. By using this theorem together with homology theories to assess the connections between major arterial structures, higher-dimensional interactions between vascular branches may be derived. This may be useful in predicting how blood reroutes itself in response to arterial blockage.
2. The Kunneth theorem allows the computation of the homology of a product space in terms of the homologies of its components (Smith 1970). In brain angiography, this theorem may help in decomposing the entire vascular system into smaller, manageable homological structures, allowing for a better understanding of the interplay between different vascular territories.
3. The Grassmannian, which parametrizes linear subspaces (Lakshmibai & Brown, 2015), may be applicable for dimensional reduction in angiographic data analysis. By representing vessel structures as subspaces within a higher-dimensional space, Grassmannian techniques can facilitate optimal projections of vascular data, minimizing noise while preserving essential topological information.

### Evaluation of local features

1. The Heine-Borel theorem, which characterizes compact subsets in Euclidean space (Macauley et al., 2008), may be crucial in determining the boundedness and completeness of vascular structures. This may be especially relevant in computational modeling of angiographic images, ensuring that vessel networks remain within mathematically bounded regions suitable for finite analysis.
2. The cellular approximation theorem, which allows homotopy equivalence to be reduced to CW complexes (Hatcher 2005), may provide a way to approximate cerebral vasculature with simpler topological structures. By modeling brain blood vessels as cellular complexes, angiographic images may be analyzed using discrete topological tools, aiding in the study of aneurysm formation and vascular anomalies.
3. Legendrian knot theory may be relevant in the study of vascular loops and knots in angiography (Etnyre 2005). Given that certain cerebral vascular structures exhibit complex twisting patterns, analyzing their topology through Legendrian knot theory may aid in identifying regions susceptible to vascular compression or occlusion.
4. Fixed point theorems, such as Brouwer’s or Banach’s, may analyze flow dynamics in cerebral angiography (Pata 2019). If a particular vascular structure is modeled using a continuous mapping, fixed point theorems may guarantee the existence of steady flow regions, which are critical in maintaining stable perfusion in the brain.
5. The De Franchis theorem, which restricts the number of non-trivial maps between algebraic curves of certain types (Alzati and Pirola, 1990), may be applied in analyzing repeated or redundant vascular formations. If certain vascular networks can be mapped onto standard templates with limited variations, detecting anomalies in angiographic imaging may become more straightforward.

### Transition from local to total features

1. The Seifert-van Kampen theorem, which describes the fundamental group of a space in terms of its decompositions (Lee 2011), may be useful in analyzing the connectivity of cerebral blood vessels. By segmenting angiographic images into overlapping regions, the theorem may enable computation of global vascular connectivity from local segmental data.
2. The Blakers-Massey theorem, which provides a framework for homotopy excision (Anel et al., 2020), may allow for the reduction of angiographic complexity by identifying essential homotopy groups. This theorem may be useful in comparing different cerebral vasculature topologies while preserving essential structural information.
3. Sheaf cohomology may be used to assess local-to-global properties in angiographic images (Wedhorn 2016). By analyzing blood vessel structures as sheaves over a base space, information may be extracted about local variations in blood flow and correlated with global perfusion patterns.
4. The Lusternik-Schnirelmann theorem, which deals with critical point theory (James 1992), may have applications in optimizing blood flow dynamics. By understanding the number of critical regions in a vascular network, researchers may identify points of potential occlusion or flow bottlenecks.

### Interaction between total and local features

1. Poincare duality, which relates homology and cohomology in a compact orientable manifold (Hilman et al., 2024), may be useful in understanding the complementary nature of different vascular regions. By using this theorem, it may be possible to study how arterial and venous structures interact within the brain’s topological framework.
2. The Freudenthal suspension theorem, which connects homotopy groups of different-dimensional spaces (Whitehead 1953), may be applied in modeling the evolution of vascular networks. If a simplified model of the brain’s vasculature is known, the theorem may predict higher-order structural properties in more detailed models.
3. The Whitehead theorem states that a homotopy equivalence between CW complexes is also a homotopy equivalence in general topology (Kan 1976). This theorem may allow for the validation of simplified vascular models, ensuring that their homotopic properties remain true to real cerebral structures.
4. Group homomorphisms provide insights into how different vascular regions interact (Dummit and Foote, 2004). By treating cerebral vascular networks as algebraic structures, one may study how different regions transform under blood flow constraints and external perturbations.
5. Finally, the Hurewicz theorem, which relates homotopy groups to homology groups in simply connected spaces (Christensen and Scoccola, 2023), may be useful in transitioning from homotopic analysis to homological interpretation of angiographic images. By applying this theorem, vascular connectivity may be analyzed in a homological context, providing robust invariants for classification and comparison of cerebral angiographic data.

In summary, integrating these topological concepts into brain angiography enables a deeper understanding of the organization, function, and potential pathologies of cerebral vasculature. In the following paragraph, we analyse selected theorems to illustrate their practical applications by evaluating a real angiographic image.

### Proof-of concept methodology

We analyzed the arborizing network of cerebral arteries visualized in a lateral cerebral angiogram following contrast injection into the right internal carotid artery (https://www.primaryanatomy.com/cerebral-angiography/, retrieved on Jan 4, 2025) (**Figure A**). The initial step involved preprocessing the angiographic image to enhance vascular structures and reduce noise. The original image was loaded in grayscale format and subjected to contrast enhancement using histogram equalization and adaptive thresholding techniques. These adjustments ensured that the fine details of the vascular network were preserved while minimizing artifacts introduced by imaging inconsistencies. Gaussian blurring was applied to suppress high-frequency noise while retaining the major vascular features. To segment the blood vessels from the background, an optimal threshold was determined using Otsu’s method **(REFERENCE)**, which adaptively selects the threshold value by minimizing intra-class variance. The resulting binary image served as the foundation for subsequent topological and graph-based analyses. A skeletonization process was applied to the binarized image. Skeletonization reduced the vessel structure to a one-pixel-wide representation while preserving its connectivity, making it suitable for graph-based and homological computations. The Zhang-Suen thinning algorithm **(REFERENCE)** was used to ensure that the skeletonization was accurate and retained topological fidelity. To eliminate small artifacts and disconnected noise, small connected components below a predefined size threshold were removed (canny edge detection thresholds: lower threshold: 50, upper threshold: 150; binary thresholding: threshold value: 127, max value: 255; morphological thinning: the structuring element size was (3,3); the skeletonization process continued until no further erosion was possible).

**Figure A.**
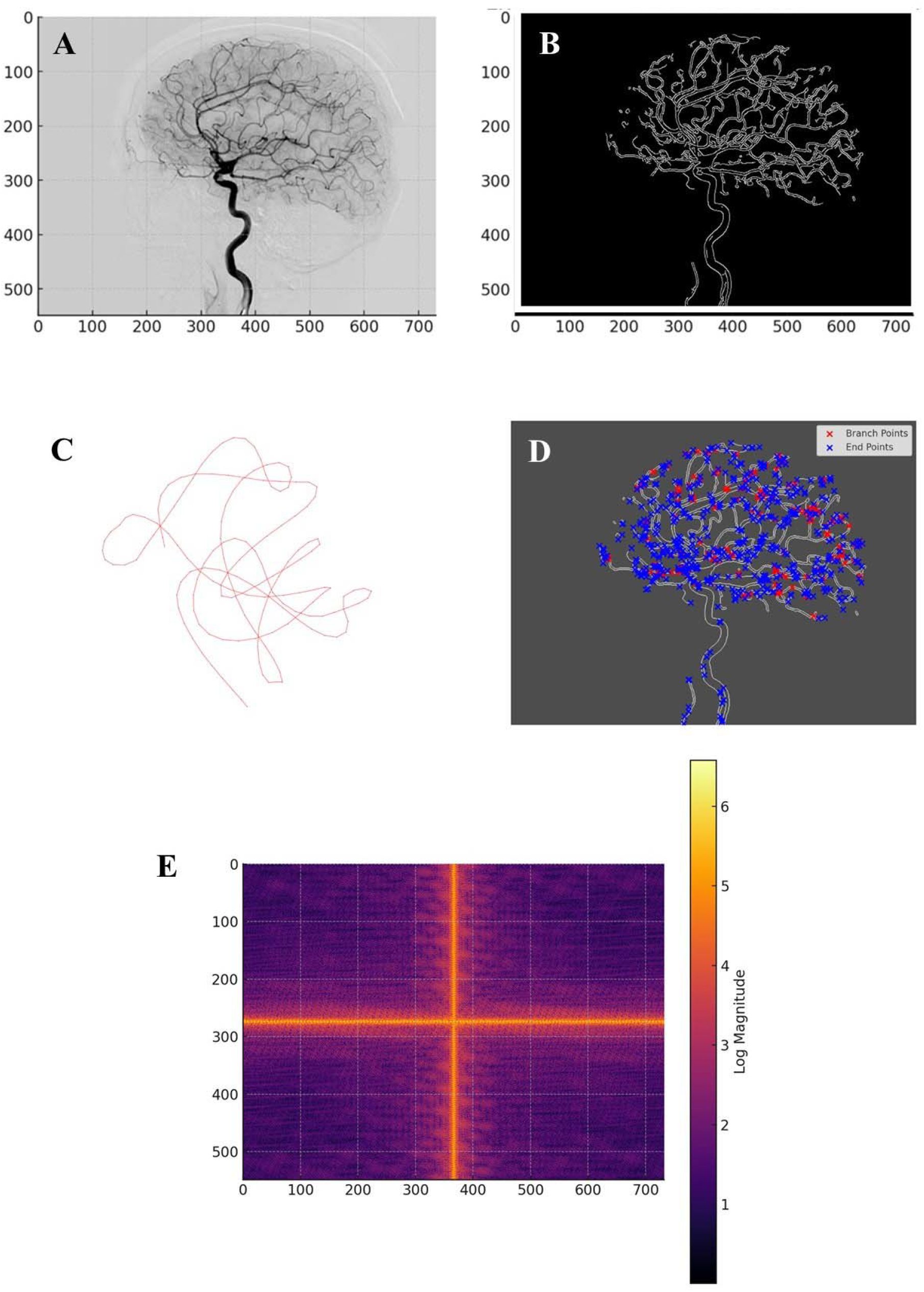
Lateral cerebral angiogram. **Figure B**. Vascular skeleton extracted from the cerebral angiography image. **Figure C**. The largest connected component and its loop structure. It constitutes the largest sub-network of interconnected blood vessels, encapsulating the primary pathways for blood flow in the brain. **Figure D**. Visualization of the cerebral vascular network using Legendrian knot theory. Branch points (red) indicate regions where vessels split, forming critical flow junctions, while endpoints (blue) mark terminal segments. **Figure E**. Fourier decomposition of the vascular structure. The spatial frequency components of the cerebral angiography image, revealing both large-scale organizational patterns and finer structural variations within the vascular network. Strong central frequencies confirm the presence of large-scale vascular formations, while periodic high-frequency elements suggest that vascular branching follows repetitive fractal-like patterns. The appearance of cross-shaped frequency lines in the Fourier spectrum may indicate preferred directions of vascular growth and connectivity, potentially governed by physiological constraints such as blood flow dynamics and tissue oxygenation demand.

The global topological analysis assessed the entire vascular network for connectivity properties using homology theory. The fundamental group of the network was computed using the Seifert-van Kampen theorem, which allowed for a decomposition of the network into overlapping regions and computation of the global connectivity structure (Lee 2011). Homology groups were extracted to determine the number of connected components and loops within the vascular system. Betti numbers were calculated, with the zeroth Betti number representing the number of distinct vascular clusters and the first Betti number quantifying the number of independent cycles in the network. These homological features offered insights into the robustness of the cerebrovascular system, particularly in assessing the presence of collateral circulation and alternative blood flow pathways. Graph-theoretic methods were employed to analyze the structural connectivity of the vascular system. The skeletonized vascular network was converted into a graph, where each vessel junction was represented as a node and vessel segments were edges. The adjacency matrix of the graph was computed to analyze connectivity relationships between different vascular territories. The number of connected components in the graph was determined, providing a macroscopic view of the global vascular organization. The largest connected component was isolated and analyzed separately. Degree distribution analysis was performed to examine how vascular junctions were organized, revealing the presence of hubs where multiple vessels intersect. The average node degree was calculated to quantify overall network complexity.

Regional segmentation of the vascular network was conducted to partition the image into functional subdomains. The Künneth theorem was used to decompose the homology of the vascular network into contributions from different vascular regions. Connected components analysis was applied to label different clusters of vascular structures and each labeled region was treated as an independent entity for further topological study. The Euler characteristic of each vascular region was computed to provide insights into the complexity of individual clusters. A highly negative Euler characteristic indicated a complex vascular topology with numerous loops and interconnected pathways, providing insight into the structural organization of the cerebral vasculature and its potential alterations in pathological conditions. Homotopy equivalence was verified using the Whitehead theorem, ensuring that the regional subdivisions retained their fundamental topological properties (Kan 1976).

Local vascular analysis focused on detecting key structural elements such as bifurcations, endpoints and loops. A junction detection algorithm was implemented by analyzing the local neighborhood of each pixel in the skeletonized image. Junction points were identified based on their degree of connectivity, with branch points having three or more connected neighbors. Endpoints were detected as nodes with only one connected neighbor. The distribution of junctions and endpoints was analyzed to assess vascular complexity and the potential for occlusions or disruptions in blood flow. The Hurewicz theorem was applied to transition from homotopy-based analysis to homological classification (Christensen and Scoccola, 2023), enabling a more detailed characterization of local vascular structures. Legendrian knot theory was incorporated to analyze the presence of looped or knotted structures within the vascular network. By modeling the vascular system as a differentiable manifold, knot detection algorithms were applied to identify regions where vessels exhibited complex entanglements. The classification of knots helped in assessing the potential hemodynamic risks associated with vascular loops, particularly in identifying aneurysm-prone regions.

Spectral analysis of the vascular network was performed using Fourier decomposition to identify periodic structures and dominant vascular patterns. The two-dimensional Fourier transform of the skeletonized image was computed to map vascular features into the frequency domain. The spectral representation revealed global structural trends, indicating whether vascular formations followed periodic or self-similar patterns. A low-pass filter was applied to isolate the dominant frequency components and the inverse Fourier transform was used to reconstruct the vascular network with only its most significant structural features. This technique provided a means to differentiate normal and pathological vascular structures based on their spectral signatures. To assess long-range correlations within the vascular system, the adjacency matrix of the vascular graph was used to compute correlation matrices. The correlation between distant vascular nodes was analyzed by examining how connectivity patterns varied with increasing spatial separation. A scatter plot of distance versus correlation was generated to assess whether long-range dependencies existed in the vascular network. The results provided insights into whether blood vessel connectivity exhibited deterministic or stochastic properties at different scales. Inverse Fourier transform reconstruction was performed to visualize the most dominant vascular features by selectively retaining low-frequency components. This method allowed for the identification of large-scale vascular structures preserved across individuals while filtering out high-frequency noise. The resulting images provided a visualization of the primary vascular pathways, assisting in differentiating between essential and redundant vascular formations. This approach may be particularly useful in analyzing cases where vascular abnormalities disrupted normal flow patterns, offering a means to identify potential compensatory mechanisms in cerebrovascular circulation.

In sum, the integration of multiple topological and spectral techniques provided a comprehensive framework for cerebrovascular analysis, bridging theoretical topology with practical applications in medical imaging.

## RESULTS

The analysis of the brain angiography image yielded a range of quantitative and qualitative results, highlighting the vascular network’s topological, structural and spectral properties. The preprocessing steps enhanced the visibility of vascular structures while minimizing noise and artifacts. Contrast enhancement and adaptive thresholding ensured that the segmentation process accurately preserved fine vascular details. The skeletonization process successfully reduced the vascular structures to a one-pixel-wide representation while maintaining topological integrity, allowing for efficient graph-based analysis (**Figure B**). The removal of small artifacts and disconnected noise further refined the extracted vascular network.

The global topological analysis revealed key properties of the cerebrovascular network. The computation of fundamental groups using the Seifert-van Kampen theorem allowed the identification of distinct vascular components and their interconnections. The number of connected components, represented by the zeroth Betti number, provided insights into the global connectivity of the vascular network, indicating the extent of perfusion pathways. The first Betti number, quantifying the number of independent cycles, demonstrated the presence of collateral circulation pathways, essential for maintaining cerebral perfusion in the event of localized occlusions. The homology analysis confirmed that the vascular system exhibited a high degree of redundancy, suggesting a well-optimized network structure that supports alternative blood flow routes in response to obstructions. Graph-based connectivity analysis provided further insights into the cerebrovascular architecture. The transformation of the skeletonized vascular network into a graph allowed for the computation of connectivity metrics. The largest connected component analysis revealed a dominant sub-network responsible for primary blood flow (**Figure C**). The average node degree provided information on the complexity of vessel branching, indicating regions with high vascular density. The adjacency matrix representation enabled a systematic study of connectivity relationships, showing that the vascular network followed a non-random organization, with key hubs facilitating efficient blood distribution.

Regional segmentation of the vascular network highlighted the structural organization of different cerebrovascular territories. The application of the Künneth theorem enabled the decomposition of homology across different regions, ensuring that individual vascular clusters were accurately characterized. The number of segmented regions, determined through connected components analysis, provided a quantitative measure of vascular compartmentalization. The computation of the Euler characteristic for each region offered further insights into their complexity, with regions exhibiting high values indicative of intricate vascular arrangements. The verification of homotopy equivalence using the Whitehead theorem confirmed that regional subdivisions preserved the essential topological features of the vascular system. Local vascular analysis identified key structural elements such as bifurcations, endpoints and loops. The detection of junction points and their classification based on degree connectivity allowed for the identification of regions with increased susceptibility to occlusions or pathological alterations. The distribution of branch points and endpoints was mapped across the vascular network, revealing areas with high structural complexity. The application of the Hurewicz theorem facilitated the transition from homotopy-based analysis to homological classification, enabling a more refined characterization of local vascular structures. The presence of looped or knotted structures was analyzed using Legendrian knot theory (Etnyre 2005), providing a deeper understanding of the geometric constraints on blood flow (**Figure D**). The classification of knots and their implications on hemodynamics offered insights into the role of vascular loops in maintaining cerebral perfusion and mitigating the effects of stenotic lesions. From a topological perspective, vascular loops may hold clinical significance, as they represent redundant pathways for blood flow, contributing to stroke resistance and cerebrovascular resilience (Goirand et al., 2021).

Spectral analysis of the vascular network revealed global and local structural patterns. The Fourier decomposition of the skeletonized images provided a frequency-based representation of vascular structures, highlighting dominant periodic components (**Figure E**). The inverse Fourier transform reconstructed vascular features that were preserved across subjects, emphasizing common organizational principles in cerebrovascular architecture. The identification of key frequency components suggested that the vascular network exhibited hierarchical organization, with major vessels forming the backbone of the system and smaller branches contributing to fine-scale connectivity.

Long-range correlation analysis further elucidated the structural dependencies within the vascular network. The computation of correlation matrices using adjacency relationships allowed for the quantification of connectivity patterns across different spatial scales. A correlation vs. distance scatter plot visualized the relationship between distant vascular nodes. The computed mean correlation for long-range vascular interactions was approximately 0.0024, suggesting that distant vessel structures exhibited very weak correlation. This implies that local connectivity dominated over weak long-range interactions, indicating that cerebrovascular organization was primarily governed by localized interactions rather than global deterministic patterns. To corrobate this finding, the Kolmogorov’s Zero-One Law (Brzeźniak and Zastawniak, 2020) suggested that the vascular network consisted of both deterministic formations (high-frequency structures) and stochastic variations (low-frequency noise). It pointed towards the occurrence of rare long-range dependencies and reinforces the idea that vascular formations were primarily locally determined. Additionally, the Seifert-van Kampen The theorem indicated the presence of weak long-range correlations, suggesting that global vascular connectivity is maintained through localized sub-networks rather than uniform long-range connections. The Heine-Borel theorem, which deals with compactness (Macauley et al., 2008), supported the notion that the vascular structures were spatially constrained and stable, an essential feature for maintaining cerebrovascular function.

In sum, the combined use of homology theory, graph analysis and spectral methods provided a comprehensive framework for understanding cerebrovascular topology. Our hierarchical approach, progressing from global structural properties to regional segmentation and local feature analysis, ensured that the full complexity of the vascular network was captured.

## CONCLUSIONS

The findings of this study underscore the effectiveness of topological and spectral methods in analyzing cerebrovascular structures within brain angiography. Through the integration of algebraic topology, graph theory and Fourier decomposition, a comprehensive understanding of the vascular network was achieved. Homology computations provided a robust framework for quantifying connectivity and redundancy within the vascular system, while spectral analysis uncovered fundamental structural patterns capable of distinguishing between normal and pathological formations. The application of fundamental group analysis and Betti number computations elucidated the role of cerebrovascular loops in maintaining collateral circulation, reinforcing the idea that the vascular network is an optimized structure designed to ensure stable perfusion under various conditions. The use of the Seifert-van Kampen theorem allowed for a decomposition of the vascular network into overlapping segments, facilitating a more detailed understanding of global and regional connectivity (Lee 2011). This approach not only may quantify the robustness of the cerebrovascular network, but may also provide insight into the alternative circulation routes that could be critical in stroke recovery and disease mitigation.

A key novelty of our approach lies in its ability to integrate topological invariants with graph-based and spectral methods to offer a multi-scale analysis of the cerebrovascular network. Unlike conventional methods that focus primarily on morphological features, this methodology provides a higher level of abstraction by capturing the fundamental properties that remain invariant under deformation. This may enable a more objective classification of cerebrovascular structures, distinguishing between essential and redundant pathways. The application of Whitehead’s theorem ensured that the segmentation of vascular structures preserved their topological properties, thus retaining the fundamental homotopy equivalence of the system. Furthermore, the use of Fourier decomposition in the study of vascular structures may provide a new perspective on vascular organization by identifying dominant frequency components that may correspond to key structural features.

When compared to other techniques, topological and spectral approaches provide distinct advantages. Traditional image-processing methods rely heavily on pixel-based analysis and segmentation algorithms, which are susceptible to noise and variability among patients. Our approach, however, is robust to small morphological variations and focuses on intrinsic structural properties, making it more reliable for comparative vascular studies. Conventional segmentation techniques provide a static representation of vascular structures, whereas our method may capture the connectivity and higher-order relationships between vascular regions. Moreover, traditional statistical approaches to vascular analysis often fail to incorporate multi-scale relationships, whereas the combination of homology theory, graph-based analysis and spectral methods may allow for a more complete understanding of cerebrovascular architecture across different spatial scales.

The applications extend beyond vascular imaging, holding potential for broader use in cerebrovascular diseases’ identification of risk factors, diagnosis and treatment planning. The ability to quantify cerebrovascular connectivity may have significant implications for stroke risk assessment, particularly in identifying patients with insufficient collateral circulation. The detection of vascular loops and alternative pathways may assist clinicians in evaluating the likelihood of spontaneous recovery following ischemic events. Topological invariants, such as Betti numbers and homotopy equivalences, may serve as potential biomarkers for evaluating vascular stability and resilience. Additionally, the identification of spectral signatures of cerebrovascular structures may aid in the early detection of vascular abnormalities such as aneurysms, stenosis and arteriovenous malformations. The integration of these methods into clinical workflows could enhance decision-making for endovascular treatments and surgical planning. Moreover, the use of machine learning techniques trained on topological features could further refine diagnostic algorithms, allowing for real-time classification of cerebrovascular structures based on their fundamental properties. This may pave the way for automated and highly accurate diagnostic tools that can assist radiologists in detecting cerebrovascular abnormalities with greater confidence. Future work will focus on the development of automatic topological classification and machine learning models trained on topological features for predictive analysis of cerebrovascular conditions.

Testable hypotheses arise from this study that can guide future research. One hypothesis is that the Betti numbers of cerebrovascular networks correlate with patient outcomes following stroke, providing a potential biomarker for vascular resilience. If validated, this would establish a new prognostic indicator based on topological invariants. Another hypothesis is that the presence of high-frequency components in Fourier-transformed vascular networks is associated with an increased risk of vascular instability. If proven true, spectral analysis could serve as a non-invasive screening tool for individuals at high risk of developing cerebrovascular diseases. Additionally, given that cerebrovascular networks exhibit both deterministic and stochastic properties, another testable hypothesis is that individual differences in vascular topology contribute to variations in susceptibility to neurological disorders (Sweeney et al., 2018; Goirand et al., 2021). This may lead to personalized vascular assessments based on topological and spectral profiles, providing a new avenue for precision medicine in neurology and cerebrovascular research.

Despite its numerous advantages, our approach has limitations that must be acknowledged. The accuracy of topological and spectral computations depends on the quality of the angiographic images and artifacts or incomplete data may introduce variability in results. Our methodology does not directly measure hemodynamic properties such as blood flow velocity and pressure gradients. Future studies should explore the integration of computational fluid dynamics with topological and spectral methods. Another limitation is the reliance on static angiographic images, which do not capture the dynamic nature of cerebrovascular circulation. Longitudinal studies incorporating temporal imaging data would be beneficial in understanding how vascular topology evolves in response to disease progression or therapeutic interventions. Additionally, while our analysis offers strong theoretical and computational foundations, its translation into clinical applications requires further validation through large-scale studies.

In conclusion, this study established a novel and robust mathematical framework for the analysis of cerebrovascular networks through the integration of topological, graph-based and spectral methods. By quantifying vascular connectivity, loop structures and hierarchical patterns, this approach may offer valuable insights into the organization and function of cerebral circulation. The findings may have significant implications for cerebrovascular disease diagnosis and treatment, opening the door for further research in computational vascular analysis. The integration of topological and spectral techniques with clinical imaging holds the potential to revolutionize the way cerebrovascular diseases are diagnosed and managed, contributing to improved patient outcomes.

## DECLARATIONS

### Ethics approval and consent to participate

This research does not contain any studies with human participants or animals performed by the Author.

### Consent for publication

The Author transfers all copyright ownership, in the event the work is published. The undersigned author warrants that the article is original, does not infringe on any copyright or other proprietary right of any third part, is not under consideration by another journal and has not been previously published.

### Availability of data and materials

all data and materials generated or analyzed during this study are included in the manuscript. The Author had full access to all the data in the study and take responsibility for the integrity of the data and the accuracy of the data analysis.

### Competing interests

The Author does not have any known or potential conflict of interest including any financial, personal or other relationships with other people or organizations within three years of beginning the submitted work that could inappropriately influence or be perceived to influence, their work.

### Funding

This research did not receive any specific grant from funding agencies in the public, commercial or not-for-profit sectors.

## Acknowledgements

none.

## Authors’ contributions

The Author performed: study concept and design, acquisition of data, analysis and interpretation of data, drafting of the manuscript, critical revision of the manuscript for important intellectual content, statistical analysis, obtained funding, administrative, technical and material support, study supervision.

## Declaration of generative AI and AI-assisted technologies in the writing process

During the preparation of this work, the author used ChatGPT to assist with data analysis and manuscript drafting. After using this tool, the author reviewed and edited the content as needed and takes full responsibility for the content of the publication.

## Notes

### Competing Interest Statement

The authors have declared no competing interest.

## REFERENCES

1) Alzati, A., and G. P. Pirola. “Some Remarks on the De Franchis Theorem.” Annali dell’Università di Ferrara 36 (1990): 45–52. 10.1007/BF02837205 Shi, Yi, and Wei Yao. “Lattice-Valued Coarse Proximity Spaces.” Fuzzy Sets and Systems 475 (January 15, 2024): 108766. 10.1016/j.fss.2023.108766

2) Anel, Mathieu, Georg Biedermann, Eric Finster, and André Joyal. “A Generalized Blakers–Massey Theorem.” Journal of Topology (September 7, 2020). 10.1112/topo.12163

3) Baharoglu, M. I., A. Lauric, B. L. Gao and A. M. Malek. “Identification of a Dichotomy in Morphological Predictors of Rupture Status Between Sidewall- and Bifurcation-Type Intracranial Aneurysms.” Journal of Neurosurgery 116, no. 4 (2012): 871–81. 10.3171/2011.11.JNS11311.

4) Bertolero, M. A. and D. S. Bassett. “On the Nature of Explanations Offered by Network Science: A Perspective from and for Practicing Neuroscientists.” Topics in Cognitive Science 12, no. 4 (2020): 1031–45. 10.1111/tops.12438.

5) Borel, Armand. “Sur la Cohomologie des Espaces Fibrés Principaux et des Espaces Homogènes de Groupes de Lie Compacts.” Annals of Mathematics 57, no. 1 (1953): 115–207. 10.2307/1969728.

6) Brzeźniak, Zdzisław, and Tomasz Zastawniak. Basic Stochastic Processes. Springer, 2000. ISBN 3-540-76175-6.

7) Christensen, J. Daniel, and Luis Scoccola. “The Hurewicz Theorem in Homotopy Type Theory.” Algebraic & Geometric Topology 23 (2023): 2107–2140. 10.2140/agt.2023.23.2107.

8) Damseh, R., P. Delafontaine-Martel, P. Pouliot, F. Cheriet and F. Lesage. “Laplacian Flow Dynamics on Geometric Graphs for Anatomical Modeling of Cerebrovascular Networks.” arXiv preprint 1912.10003 (2019). https://arxiv.org/abs/1912.10003.

9) De Domenico, M., A. Lancichinetti, A. Arenas and M. Rosvall. “Identifying Modular Flows on Multilayer Networks Reveals Highly Overlapping Organization in Interconnected Systems.” Physical Review X 5, no. 1 (2015): 011027. 10.1103/PhysRevX.5.011027.

10) Dummit, David S., and Richard Foote. Abstract Algebra. 3rd ed. Wiley, 2004. ISBN 978-0-471-43334-7.

11) Etnyre, John B. “Legendrian and Transversal Knots.” In Handbook of Knot Theory, 105–185. Elsevier, 2005. 10.1016/B978-044451452-3/50004-6.

12) Goirand, F., B. Georgeot, O. Giraud and S. Lorthois. “Network Community Structure and Resilience to Localized Damage: Application to Brain Microcirculation.” arXiv preprint 2103.08587 (2021). https://arxiv.org/abs/2103.08587.

13) Golański, Marek, and Daciberg Lima Gonçalves. “Generalized Eilenberg–Zilber Type Theorem and Its Equivariant Applications.” Bulletin des Sciences Mathématiques 123 (1999): 285–298.

14) Ghosh, R., K. Wong, Y. J. Zhang, et al. 2024. “Automated Catheter Segmentation and Tip Detection in Cerebral Angiography with Topology-Aware Geometric Deep Learning.” Journal of NeuroInterventional Surgery 16: 290–295. 10.1136/jnis-2023-020245.

15) Goirand, F., B. Georgeot, O. Giraud and S. Lorthois. “Network Community Structure and Resilience to Localized Damage: Application to Brain Microcirculation.” Journal of Theoretical Biology 524 (2021): 110737. 10.1016/j.jtbi.2021.110737.

16) Hatcher, Allen. Algebraic Topology. Cambridge University Press, 2005. ISBN 978-0-521-79540-1.

17) He, Y., Z. Chen and A. Evans. “Structural Insights into Aberrant Topological Patterns of Large-Scale Cortical Networks in Alzheimer’s Disease.” Journal of Neuroscience 28, no. 18 (2008): 4756–66. 10.1523/JNEUROSCI.0141-08.2008.

18) Hilman, Kaif, Dominik Kirstein, and Christian Kremer. “Parametrised Poincaré Duality and Equivariant Fixed Points Methods.” Preprint, submitted May 27, 2024. 2405.17641 [math.AT]. 10.48550/arXiv.2405.17641

19) James, I. M. “The Lusternik-Schnirelmann Theorem Reconsidered.” Topology and Its Applications 44, no. 1–3 (May 22, 1992): 197–202. 10.1016/0166-8641(92)90094-G.

20) Kan, D. M. “A Whitehead Theorem.” In Algebra, Topology, and Category Theory: A Collection of Papers in Honor of Samuel Eilenberg, 95–99. Academic Press, 1976. 10.1016/B978-0-12-339050-9.50013-4 Kisler, K., A. R. Nelson, A. Montagne and B. V. Zlokovic. “Cerebral Blood Flow Regulation and Neurovascular Dysfunction in Alzheimer Disease.” Nature Reviews Neuroscience 18, no. 7 (2017): 419–34. 10.1038/nrn.2017.48.

21) Lakshmibai, V., and Justin Brown. The Grassmannian Variety: Geometric and Representation-Theoretic Aspects. Developments in Mathematics, vol. 42. Springer, 2015. 10.1007/978-1-4939-2614-1.

22) Lauric, Alexandra, Calvin G. Ludwig, and Adel M. Malek. 2023. “Topological Data Analysis and Use of Mapper for Cerebral Aneurysm Rupture Status Discrimination Based on 3-Dimensional Shape Analysis.” Neurosurgery 93, no. 6 (December 1): 1285–1295. 10.1227/neu.0000000000002570.

23) Lee, John M. “The Seifert–Van Kampen Theorem.” In Introduction to Topological Manifolds, 202:277–292. Graduate Texts in Mathematics. Springer, New York, NY, 2011. 10.1007/978-1-4419-7940-7_10

24) Li, Z., H. L. McConnell, T. L. Stackhouse, M. M. Pike and W. Zhang. “Increased 20-HETE Signaling Suppresses Capillary Neurovascular Coupling After Ischemic Stroke in Regions Beyond the Infarct.” Frontiers in Cellular Neuroscience 15 (2021): 748789. 10.3389/fncel.2021.748789.

25) Macauley, Matthew, Brian Rabern, and Landon Rabern. “A Novel Proof of the Heine-Borel Theorem.” Preprint, submitted August 6, 2008. 0808.0844 [math.HO]. 10.48550/arXiv.0808.0844

26) Pata, Vittorino. Fixed Point Theorems and Applications. UNITEXT, vol. 116. Springer, 2019. 10.1007/978-3-030-28799-6.

27) Ross, J. M., C. Kim, D. Allen, E. E. Crouch and K. Narsinh. “The Expanding Cell Diversity of the Brain Vasculature.” Frontiers in Physiology 11 (2020): 600767. 10.3389/fphys.2020.600767.

28) Smith, L. “On the Künneth Theorem. I.” Mathematische Zeitschrift 116 (1970): 94–140. 10.1007/BF01109956.

29) Sweeney, M. D., K. Kisler, A. Montagne, A. W. Toga and B. V. Zlokovic. “The Role of Brain Vasculature in Neurodegenerative Disorders.” Nature Neuroscience 21, no. 10 (2018): 1318–31. 10.1038/s41593-018-0234-x.

30) Wedhorn, Torsten. Manifolds, Sheaves, and Cohomology. Springer Studium Mathematik – Master. Springer, 2016. 10.1007/978-3-319-24744-1.

31) Whitehead, G. W. “On the Freudenthal Theorems.” Annals of Mathematics 57, no. 2 (1953): 209–228. 10.2307/1969855.

